# SARS-CoV2 (COVID-19) Structural/Evolution Dynamicome: Insights into functional evolution and human genomics

**DOI:** 10.1101/2020.05.15.098616

**Authors:** Ruchir Gupta, Jacob Charron, Cynthia L Stenger, Jared Painter, Hunter Steward, Taylor W Cook, William Faber, Austin Frisch, Eric Lind, Jacob Bauss, Xiaopeng Li, Olivia Sirpilla, Xavier Soehnlen, Adam Underwood, David Hinds, Michele Morris, Neil Lamb, Joseph A Carcillo, Caleb Bupp, Bruce D Uhal, Surender Rajasekaran, Jeremy W Prokop

**Author notes:** **Corresponding Author**: Jeremy W Prokop, Address: 400 Monroe Ave NW, Grand Rapids, MI 49503 USA.

## Abstract

The SARS-CoV-2 pandemic, starting in 2019, has challenged the speed at which labs perform science, ranging from discoveries of the viral composition to handling health outcomes in humans. The small ~30kb single-stranded RNA genome of Coronaviruses makes them adept at cross species spread and drift, increasing their probability to cause pandemics. However, this small genome also allows for a robust understanding of all proteins coded by the virus. We employed protein modeling, molecular dynamic simulations, evolutionary mapping, and 3D printing to gain a full proteome and dynamicome understanding of SARS-CoV-2. The Viral Integrated Structural Evolution Dynamic Database (VIStEDD) has been established (prokoplab.com/vistedd), opening future discoveries and educational usage. In this paper, we highlight VIStEDD usage for nsp6, Nucleocapsid (N), and Spike (S) surface glycoprotein. For both nsp6 and N we reveal highly conserved surface amino acids that likely drive protein-protein interactions. In characterizing viral S protein, we have developed a quantitative dynamics cross correlation matrix insight into interaction with the ACE2/SLC6A19 dimer complex. From this quantitative matrix, we elucidated 47 potential functional missense variants from population genomic databases within ACE2/SLC6A19/TMPRSS2, warranting genomic enrichment analyses in SARS-CoV-2 patients. Moreover, these variants have ultralow frequency, but can exist as hemizygous in males for ACE2, which falls on the X-chromosome. Two noncoding variants (rs4646118 and rs143185769) found in ~9% of African descent individuals for ACE2 may regulate expression and be related to increased susceptibility of African Americans to SARS-CoV-2. This powerful database of SARS-CoV-2 can aid in research progress in the ongoing pandemic.

## Introduction

The current SARS-CoV-2 outbreak has become a global pandemic. There is an urgent need to understand the proteins coded by SARS-CoV-2 and how they can be targeted for intervention. Coronaviruses belong to the *Orthocoronavirinae* subfamily, which lies under the *Coronaviridae* family. Their 26-33kb genome consists of positive-sense, single-stranded RNA, coding for non-structural and structural proteins. To date, seven coronaviruses have been discovered that are capable of human-to-human transmission. Four of these cause the common cold (HKU1, NL63, OC43, 229E), while the other three (MERS-CoV, SARS-CoV, SARS-CoV-2) can cause more severe respiratory illnesses resulting in multi-system organ failure and death (Kasmi et al., 2020). SARS-CoV-2 shares 79% genomic similarity with SARS-CoV, linking it to the bat and human SARS-CoVs annotation. Both SARS-CoV and SARS-CoV-2 bind to the ACE2 receptor through the Spike (S) protein (Kuba et al., 2005; Yan et al., 2020). The ubiquitous presence of Coronaviridae in many animals and its relatively small genome makes these an ideal infective agent as it adapts and evolves into a highly effective pathogen. In the case of the SARS-CoV-2 genome, 96.2% of the genome is shared with a Bat coronavirus, suggesting a zoonotic origin (Andersen et al., 2020; Ceraolo and Giorgi, 2020). Initial disease propagation was detected in Wuhan, China, a major transportation hub with over 11 million people. The population density and the heavy traffic into and out of the city made for a large outbreak that spread quickly throughout the world (Dong et al., 2020; Guarner, 2020). This combination has given rise to a once in a generation pandemic.

Insights can be gathered from SARS-CoV and MERS-CoV regarding SARS-CoV-2 illness severity. Studies in mouse models have led to speculation that SARS-CoV and MERS-CoV infections cause a delayed type I IFN response, which allows for early uncontrolled viral replication. This leads to an influx of neutrophils and monocytes/macrophages resulting in a hyperproduction of cytokines causing pneumonia, acute respiratory distress syndrome (ARDS), and global sepsis. In SARS-CoV-2, patients experience similar changes in neutrophils and lymphocytes, indicating that SARS-CoV-2 infection severity may closely depend on a delayed response of the innate immune system (Prompetchara et al., 2020). Early Chinese data provides remarkable insights into how SARS-CoV-2 drives lethality through a sepsis driven multiple organ failure model (Zhou et al., 2020), including early spike in ferritin, cytokine storm, and injury to the cardiac system (Mehta et al., 2020; Shi et al., 2020). Understanding of the SARS-CoV-2 genome could provide critical insights into the complex interplay with the host genome driving disease progression in severe risk group patients, allowing for the identification of potential target sites for intervention.

Several *Coronaviridae* proteins have been highly studied as targets of intervention to prevent infection spread. One of these proteins, the N protein is of particular interest as it interacts with host ribosomal subunits and has been shown to suppress non-sense mediated decay (NMD) of viral mRNA by the host cell (Malle, 2020, p. 2; Wada et al., 2018). Enzymes encoded by SARS-CoV-2 such as the 3-chymotrypsin (3C)-like protease, RNA-dependent RNA polymerase, and papain-like protease are potential targets of drugs (Wu et al., 2020). Proteins of the virus including N, nsp9, nsp13, nsp15, ORF3a and ORF6 have been shown to target innate immune signaling pathways (Malle, 2020). The S protein, surface expressed, enters cells through ACE2/SLC6A19 membrane receptor contact similarly to SARS-CoV, but with higher binding affinity (Yan et al., 2020). This interaction has the potential to be therapeutically inhibited through antibody neutralization (Walls et al., 2020). ACE2 is highly expressed in the heart, small intestine, kidney, thyroid, breast, arterial walls, adipose and testis with lower number of cells in lung, oral/nasal cavity, pancreas and liver (Hamming et al., 2004, p. 2). Nasal epithelial cells play a critical role in SARS-CoV-2 (Sungnak et al., 2020). In the lung, ACE2 is expressed in the type 2 pneumocytes, putative progenitor cells of alveolar epithelia linked to lung fibrosis pathways (Barkauskas et al., 2013; Uhal et al., 2013). The Spike-ACE2 complex is further processed by TMPRSS2 for internalization of the virus, which has been postulated to be a target site with protease inhibitors (Hoffmann et al., 2020). Currently, it is not well understood how the Spike-ACE2 complex is impacted by viral or human variants, suggesting a need for further research.

As we learn more about viral pathogenesis, hopefully the current outbreak will be brought under control and future outbreaks prevented. In this current work we developed the Viral Integrated Structural Evolution Dynamic Database (VIStEDD) for SARS-CoV-2 proteome, enriching VIStEDD using evolutionary insights of viruses and human variant mapping for potential functional outcomes.

## Results

### SARS-CoV-2 dynamicome database

A total of 24 proteins of SARS-CoV-2 (Table 1) were run through our standardized workflow (Figure 1) consisting of protein structure assessment, setting of protein protonation at pH 7.4, energy minimization in water with NaCl, 20 nanoseconds of molecular dynamic simulations, analysis of the movement trajectories, sequence identification from the nr database, mapping of conservation onto structure/dynamics, and assessment of interactions with known binding partners. We extracted 55,390 sequences homologous to the SARS-CoV-2 proteins that consist of 9,701 amino acids (Table 1). The uneven sequence depth for each of the proteins made it necessary to utilize a z-score conservation calculation for each of the amino acids in the proteins so as to normalize, using the average (z-score of 0) as the starting point (yellow), followed by z-scores of 0-0.5 = yellow, 0.5-1 = bright orange, 1-1.5 = orange, 1.5-2 = dark orange, >2 = red (Figure 2). The dynamics and evolution for each protein were integrated together into the Viral Integrated Structural Evolution Dynamic Database (VIStEDD) available at prokoplab.com/vistedd. VIStEDD has been built for the addition of future viruses. Within the SARS-CoV-2 page is a list of each of the 24 proteins in the format of Table 1, where each protein can be clicked to assess data. On the page for each protein is a link to the individual protein data folder system, a video of the protein rotating with conservation, details of the protein function, a widget to purchase a 3D print of the protein at cost of production, the amino acid movement from molecular dynamic simulations (mds), and the table of data for each amino acid of the protein. If protein interactions structures are known, information is present with a link to PPI data. For example within the nsp10 data (prokoplab.com/nsp10/), structures of nsp10 interacting with either nsp14 or nsp16 are available, both of which show highly conserved contact sites of interaction. As we continue to advance VIStEDD, we anticipate the addition of more material within each page.

**Table 1.**
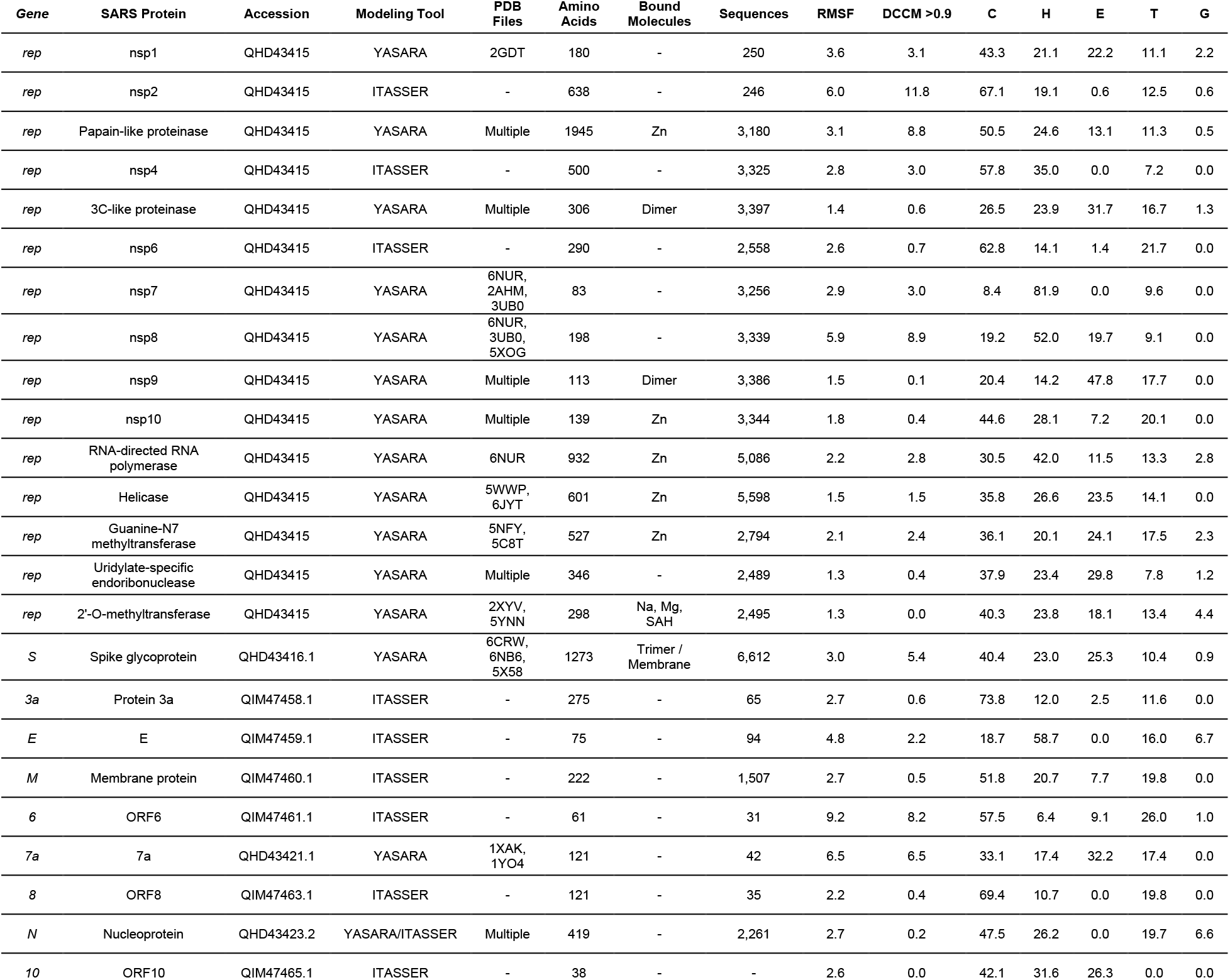
SARS-CoV2 proteins analyzed. Shown for each protein includes the gene, accession #, tool used to model the protein, known PDB files used for modeling, the number of amino acids in the protein, bound molecules, the number of sequences for evolution, the average root-mean squared fluctuation (RMSF) per residue, the average number of amino acids correlated with each amino acid greater than 0.9 (DCCM), and the percent of each proteins secondary structure (C-coil, H-helix, E-beta sheet, T-turn, G-3 turn helix).

**Figure 1.**
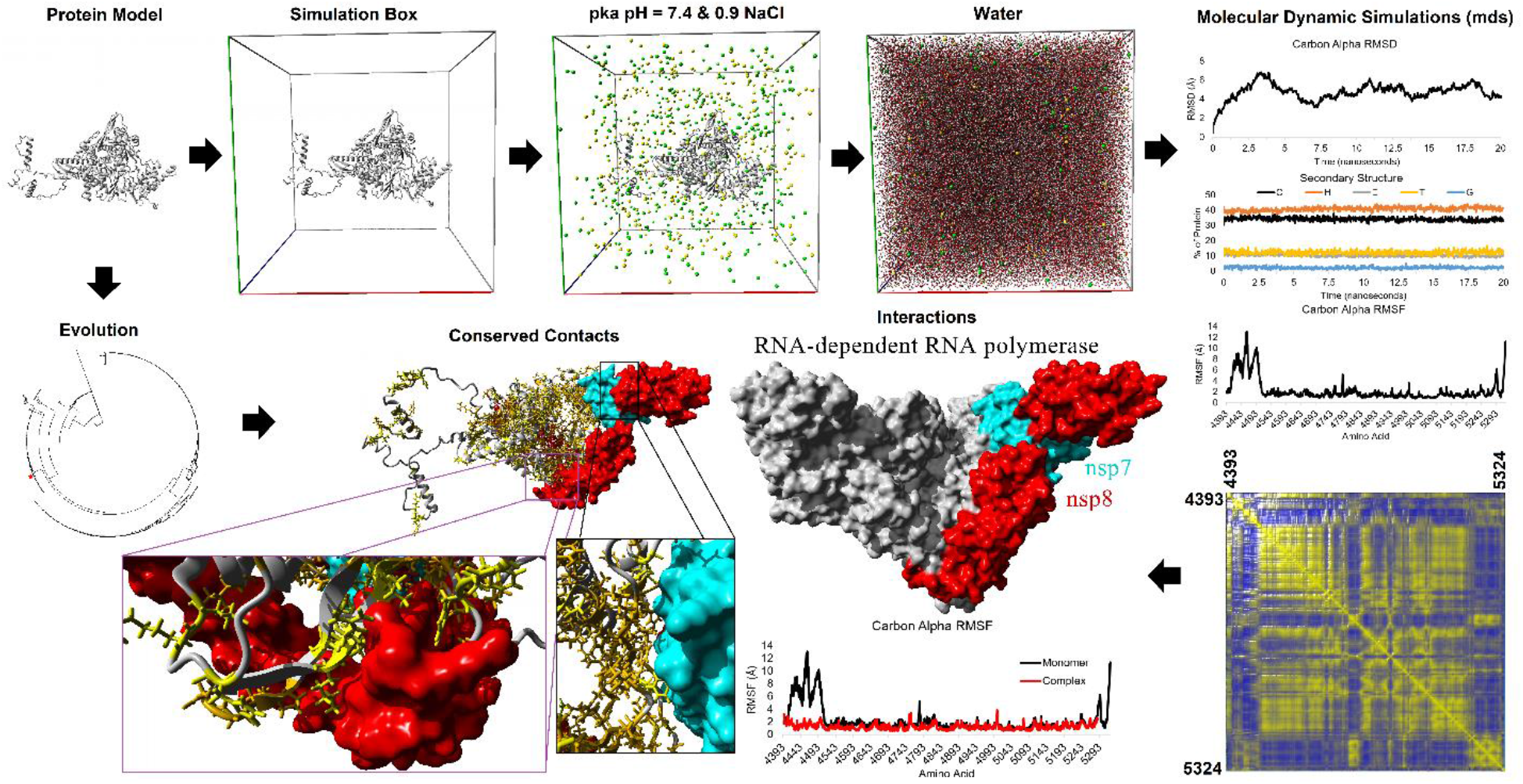
SARS-CoV2 structural/evolution dynamicome workflow. Shown is data for the RNA-directed RNA polymerase.

**Figure 2.**
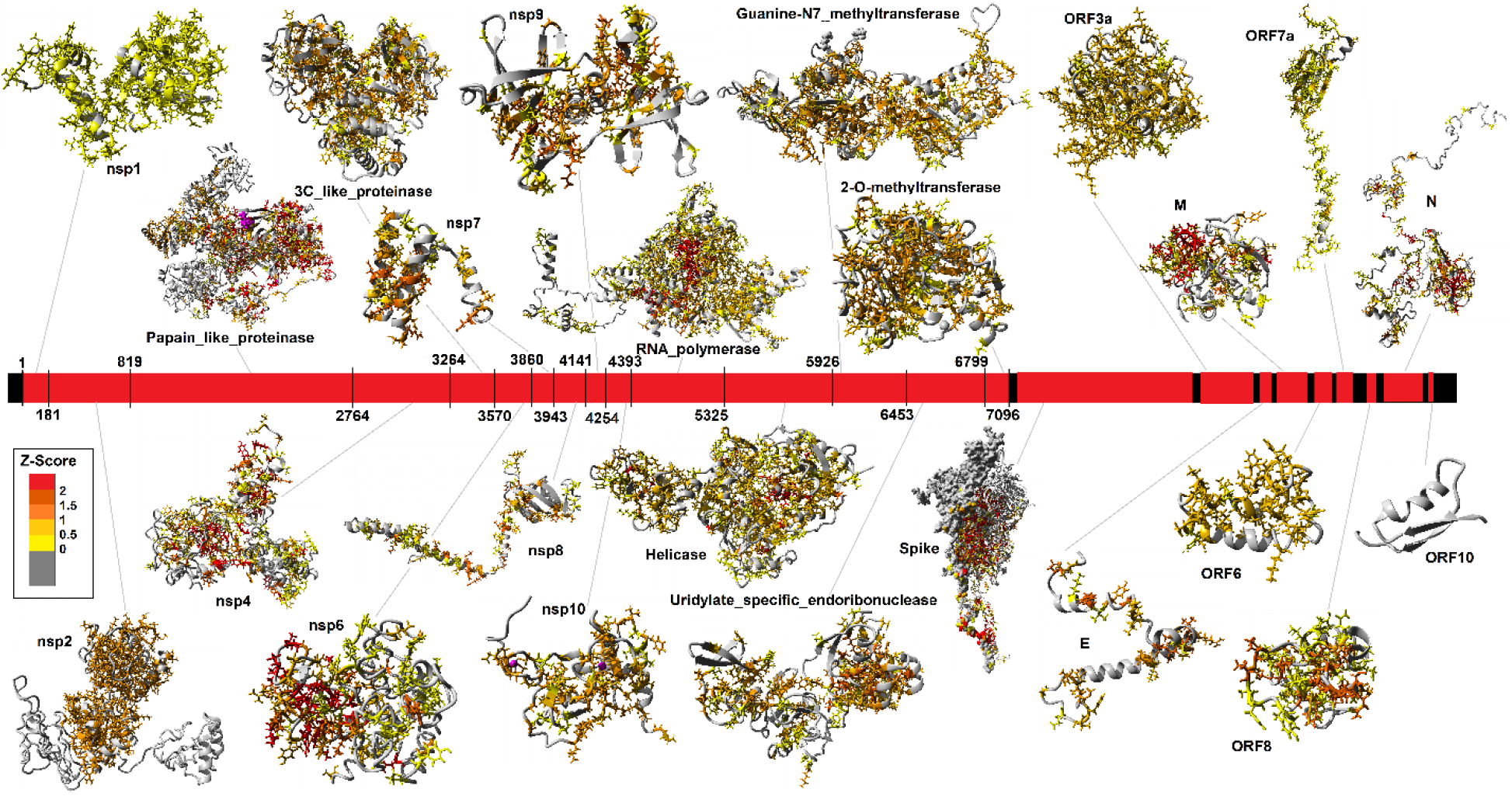
SARS-CoV2 structural/evolution models. Shown in the middle is the viral RNA with protein coding genes in red. Gray lines connect the RNA region to each protein. Colors on the models are based on Z-score levels of conservation for each protein based on extracted sequences (Table 1). Gray amino acids fall below the average conservation (value <0) for each protein, yellow = 0-0.5, light orange = 0.5-1, orange = 1-1.5, dark orange = 1.5-2, red = >2.

The raw data of each protein is the strength of VIStEDD (drive.google.com/drive/folders/1dXBJpLo3bay1JQ9BckUsVcTViv6P0w1q?usp=sharing). For each protein present in VIStEDD, we have generated in the root folder of the protein a fasta sequence file, PDB file of the protein structure, protein models with conservation (sce = YASARA scene, pse-PyMOL scene), high-resolution image of conservation, molecular video of the conservation rotating around the y-axis (mpg and mp4), and compiled conservation and dynamics data for each amino acid (csv or tab delineated). Five folders of data are also present: 1) 3D-containing a vrml 3D printing file (also zipped) of conservation mapped on the protein, 2) genomics-contains aligned reads of the species sequences extracted, 3) mds-contains all of the trajectory files for the mds, 4) Report-containing all of the analysis files from YASARA assessment of mds, and 5) tab-containing all of the tab delineated analysis files of the mds. All the 3D files can be ordered from Shapeways with the web links provided in Table S1. Another folder (PPI) contains data for all of the mds performed on protein-protein interactions including Spike-ACE2-SLC6A19, TMPRSS2, the polymerase complex, the N complex, and ACE2_S_Database consisting of 235 species ACE2 sequences modeled with Spike interaction, energy minimized, and binding or potential energy calculated. A tab delineated file is in the ACE2_S_Database to label the species for each numbered complex.

From the mds of all proteins, a total of 6,594,981 atoms including water, there is an average movement per amino acid (root-mean squared fluctuation, RMSF) of 3.2Å with 3 amino acids correlated per residue >0.9 based on dynamics cross correlation calculation. The secondary structures of the proteins are relatively similar from the beginning of the simulation compared to the end (Figure 3A), with the largest percent coiled (C, 45.02%) followed by helix (H, 25.42%), beta-sheet (E, 15.08%), turns (T, 13.44%), and 3 turn helix (G, 1.05%). On their own, before protein-protein interactions (PPI) the nsp7, nsp8, nsp9, E, 3C-proteinase, and RNA-directed RNA polymerase are the most helical and beta-sheet containing proteins while Protein 3a, ORF8, nsp2, nsp6, and nsp4 are the most disordered with coil composition (Figure 3B). Several of the proteins contain high movement of the structure (ORF6, 7a, nsp8, and nsp2) and two of the proteins (Papain-like proteinase and Spike glycoprotein) have more overall lower movement indicative of hydrophobic collapse, with a high number of correlated amino acids per residue (Figure 3C).

**Figure 3.**
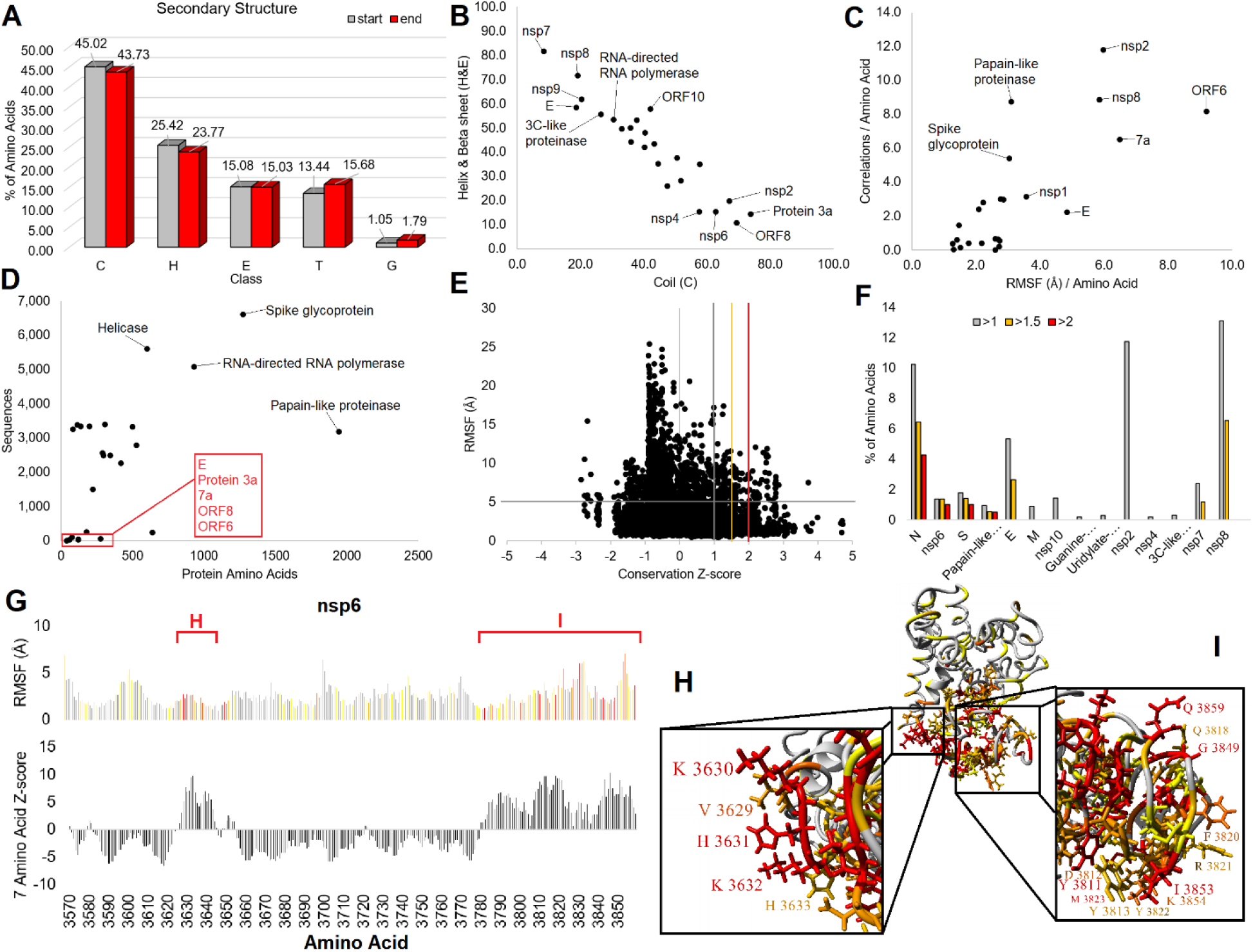
SARS-CoV2 structural/evolution statistics. **A)** The percent secondary structure for all proteins at the start of molecular dynamic simulations (gray) and at the end (red). Classes consist of C-coil, H-helix, E-beta sheet, T-turn, G-3 turn helix. **B)** Breakdown for each protein for secondary structure percent that is coil (x-axis) vs. helix/beta-sheet (y-axis). The top on each end are labeled. **C)** Breakdown of molecular dynamic simulation data for each protein showing the average amino acid root-mean squared fluctuation (RMSF, Å, x-axis) vs. the average number of correlated amino acids per residue (y-axis). Proteins with high movement are labeled. **D)** Plot of the number of amino acids in each protein vs the number of BLAST extracted sequences. Labeled are those that have high numbers of identified sequences (black) and those with only a few (red). **E)** The conservation Z-score (x-axis) vs. the RMSF (y-axis) for all amino acids analyzed in SARS-CoV2. The lines represent cutoffs used for panel F, with those >5Å for RMSF and 1-1.5 (gray), 1.5-2 (orange), or >2 (red) value for Z-score cutoffs. **F)** The % of each proteins amino acids that fall into identified groups from panel E, representing identified highly dynamic and conserved amino acids. **G)** Conservation/dynamics of nsp6 amino acids. Shown on the top is the RMSF of each amino acid of nsp6 with colors corresponding to cutoffs of panels E-F. Shown on the bottom is a sliding window calculation of 7 amino acids for additive z-scores to map two highly conserved sites (shown in panels H-I). **H-I)** Protein model of nsp6 with z-score coloring of Figure 2. Shown are the two sites of high conservation with amino acids labeled.

The largest proteins of SARS-CoV-2 contain the highest number of sequences extracted for homology including the Papain-like proteinase, RNA-directed RNA polymerase, Spike glycoprotein, and the Helicase (Figure 3D). Several of the proteins including E, protein 3a, 7a, ORF8, and ORF6 have a low number of mapped sequences (Figure 3D), while ORF10 has no other identified sequences. Plotting the conservation relative to the mds based movement, RMSF, for each of the 9,701 amino acids of SARS-CoV-2 can be used to identify critical sites of proteins and under high selection that might be targeted (Figure 3E). Highly dynamic and conserved amino acids are the prime location for critical PPI. Therefore, we mapped sites of high movement, >5Å, with conservation 1-1.5 (gray), 1.5-2 (orange), or >2 (red) standard deviations higher than the mean of the protein. Clustering these sites to the % of amino acid reveals a likely high selection of dynamic amino acids (Figure 3F). The nsp2 protein with low coverage of species has some suggested PPI contacts conserved in the range of 1-1.5 standard deviations conservation. Proteins nsp7 and nsp8, which are known to contact the RNA-dependent RNA polymerase (Figure 1), have several amino acids conserved in the 1.5-2 standard deviation range with high dynamics, as the mds of PDB 6m71 and 7btf show stability and correlation to PPI within VIStEDD.

### nsp6 and conserved

The Papain-like protease and nsp6 have conserved sites >2 standard deviations (Figure 3F). The SARS-CoV-2 nsp6 protein is known to interact with multiple ATPases of vesicle trafficking (Gordon et al., 2020) and interacts with nsp3 and nsp4 to induce double-membrane vesicles (Angelini et al., 2013). Nsp6 protein also interacts with the Sigma receptor, which is thought to regulate ER stress response (Gordon et al., 2020) and blocks ER-induced autophagosome/autolysosome vesicle that restricts viral production leading to the generation of small autophagosome vesicles, thereby limiting their expansion (Cottam et al., 2014). To date, no structures of nsp6 have been solved even though the protein is present in 2,558 *Coronaviridae* genomes. We identify two regions with minimal conserved hydrophobic collapse (Figure 3G) consisting of mostly coiled secondary structure (Figure 3B). These two regions of conservation (Figure 3G) cluster together with multiple charged and aromatic amino acids that would tend to drive PPI (Figure 3 H-I). Moving forward these nsp6 sites could be critical regions to target with therapeutics for the broad *Coronaviridae* proteins.

### Nucleocapsid (N) data insights

Both the S and N proteins have multiple sites conserved >2 standard deviations with high dynamics (Figure 3F), with N having around 4% of its amino acids falling into this category. These two proteins are further dissected below. The SARS-CoV-2 N protein has been shown to interact with multiple RNA processing and stress granule proteins (Gordon et al., 2020). Using 2,261 sequences (Figure 4A) we mapped 4 highly conserved regions of N (Figure 4B). The conservation of region 1 contributes to a highly conserved hydrophobic, aromatic core consisting of several beta strands (Figure 4C-D). Conserved region 3 consists of several serine amino acids that are likely phosphorylated and a potential 14-3-3 binding motif (Figure 4D). Molecular dynamics of N suggests three regions of structural organization, with region 1 corresponding to a domain with structural folding (Figure 4E). Amino acids 331-333 were the most conserved yet dynamic site of N (Figure 4F). The C-terminal region of N is known to form a multimer complex (Figure 4G) with amino acids 331-333 fitting into contacts of the subunits (Figure 4H). From the multimer N structure, conserved amino acids V270, F274, R277, N285, G287, F286, and D288 are clustered and surface exposed (Figure 4I). These sites are likely to contribute to PPI and warrant future investigations.

**Figure 4.**
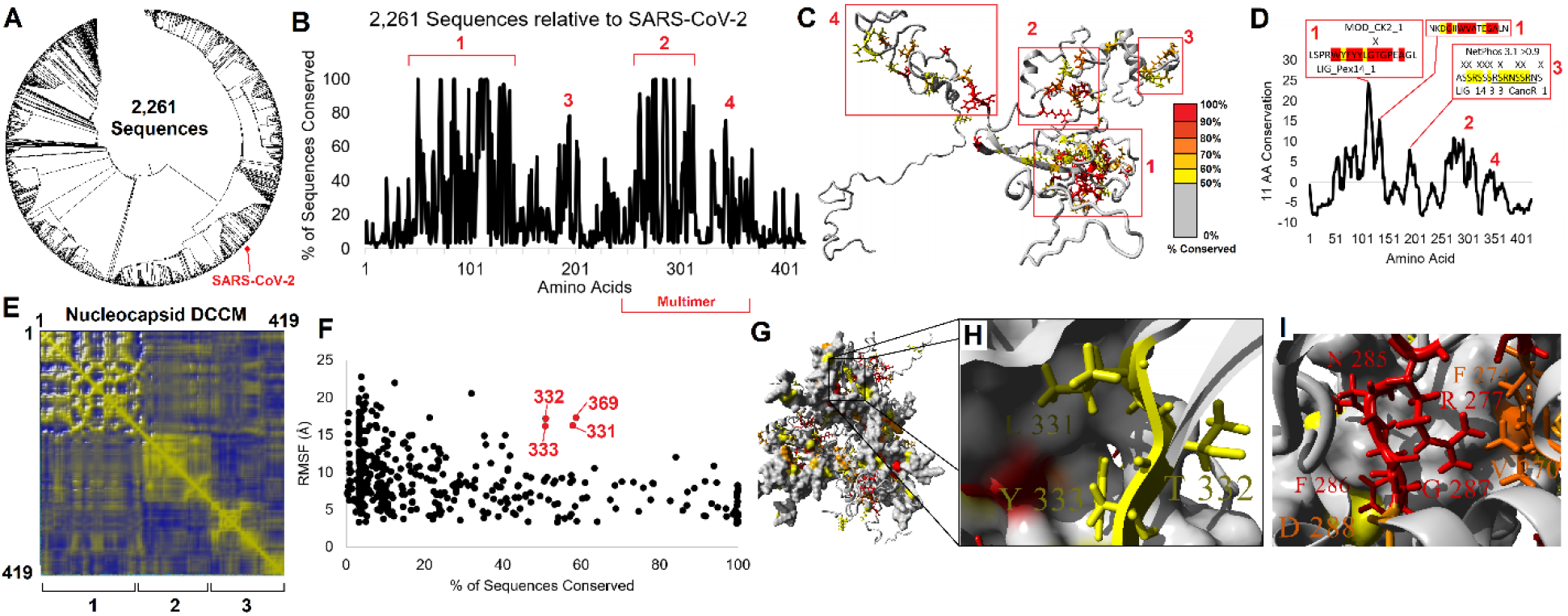
SARS-CoV2 protein N (Nucleocapsid) conserved dynamic amino acids. **A**) Phylogenetic tree of 2,261 sequences extracted from the nonredundant (nr) protein sequence database for N. **B**) Conservation of amino acids based on sequences from panel A. Four regions of conserved function are identified in red. **C)** Model of N with conservation (% of 2,261 sequences) colored (gray=<50%, yellow=50-60%, light orange = 60-70%, orange =70-80%, dark orange = 80-90%, red = >90%. The four regions of panel B are boxed and labeled in red. **D)** Top conserved motifs in N. The Z-scores for N conservation were placed on an 11 codon sliding window to identify regions of interest in the four conserved regions of panel B. **E**) Dynamics cross-correlation matrix (DCCM) of amino acids on the nucleocapsid protein assessed with molecular dynamic simulations. **F**) Intersection of conservation and dynamics of the protein with highly dynamic and conserved amino acids in red. **G)** Multimeric model of N with coloring based on panel E. **H)** Zoom in view of black box from panel F showing amino acids 331-333 identified in panel D. **I)** Conserved region 2 from panel B identifying top amino acids on the multimer model.

### Posttranslational modification analysis

Building on our insights for N, we expanded to a systematic analysis of posttranslational modifications for SARS-CoV-2 proteins that was integrated into our amino acid matrix insights (Figure 5). Current literature on SARS-CoV-2 modifications focus on conserved and novel glycosylation and phosphorylation sites of the S protein (Xiao et al., 2003). With host-pathogen interaction being regulated by modifications, it is now a target for pharmacotherapy. Data from SARS-CoV suggest modifications in the N, M, E, 3a, nsp4, and nsp9 proteins that have yet to be explored in SAR-CoV-2 (Fung and Liu, 2018; Nal et al., 2005; Oostra et al., 2006). Specifically, the N protein was shown to undergo extensive acetylation, phosphorylation, sumoylation, cleavage, and ADP-ribosylation (Fan et al., 2006; Surjit et al., 2005). Inhibition of several cellular kinases (CK2 and CDK) were shown to impact proper localization of the N protein, trapping it within the nucleus of the host cell, further highlighting the need for N protein phosphorylation to carry out proper function (Surjit and Lal, 2008). With these PTMs playing a vital role in proper virion assembly, they must be further explored and understood in SARS-CoV-2 to elucidate all options for targeted pharmacotherapy.

**Figure 5.**
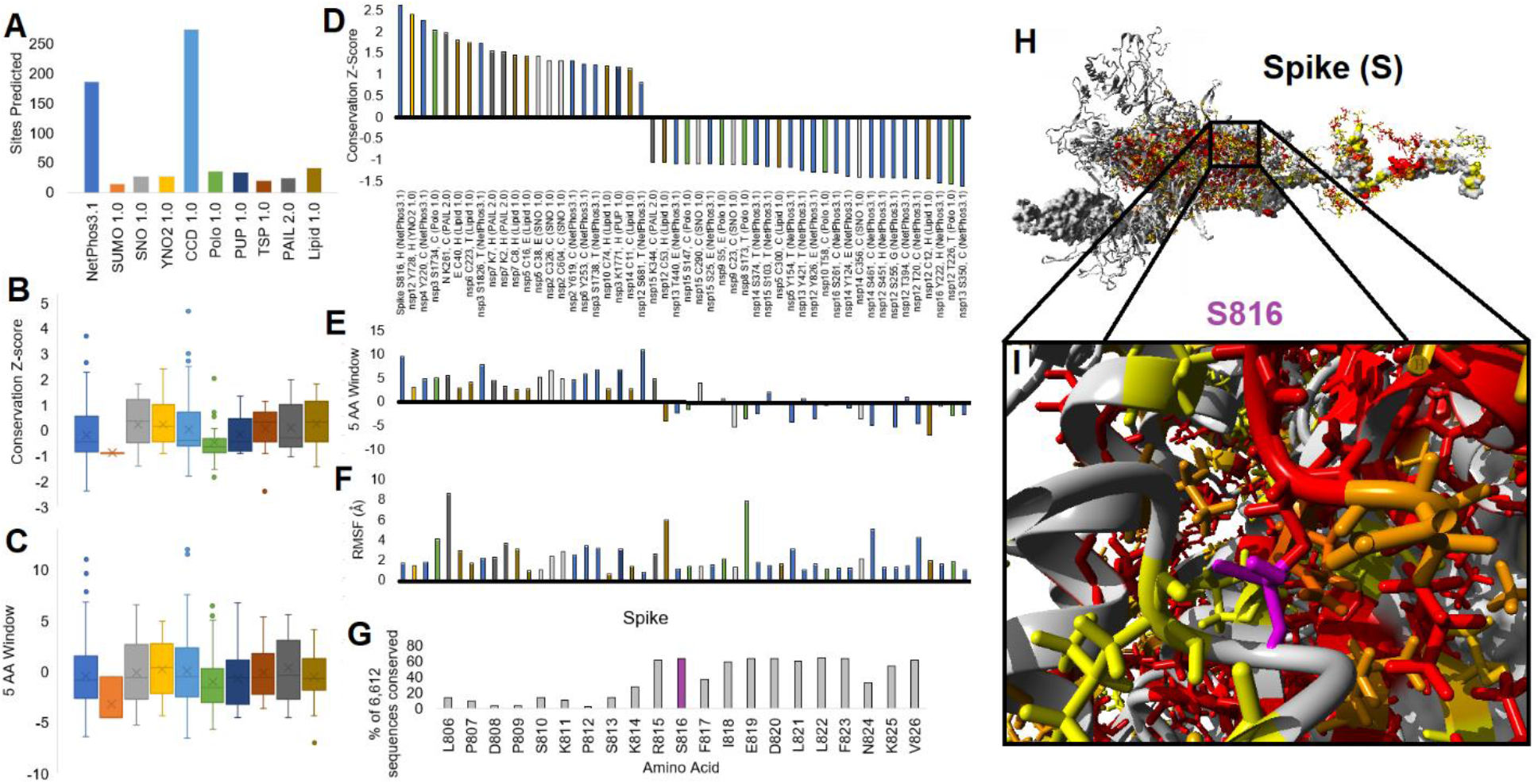
Posttranslational modification screening of SARS-CoV-2. **A**) Ranking of functional predicted sites using 10 different tools. **B-C**) Conservation Z-score (**B**) and Z-Scores put on a five codon sliding window for additive motif conservation (**C**) of each site from panel A for each tool shown as a box and whisker plot. **D-F**) Top conserved sites (left) and least conserved sites that are unique to SARS-CoV-2 (right) for functional predictions. Colors of bars correspond to tools of panel A. Each functional amino acid is labeled as Protein Amino Acid, secondary structure (tool). Data is shown for the Z-Score of conservation (**D**), the 5 amino acid sliding window (**E**), and structural movement (**F**). G) The highest conserved site throughout the database found on Spike at S816 with conservation of amino acids around the site. **H-I**) Highlighting the phosphorylation prediction (magenta) on Spike for S816 (**H**) with zoom in view of site on one of the monomers (**I**).

With high filters on each tool we identify 186 NetPhos3.1 (phosphorylation), 15 SUMO1.0 (Sumo binding or SUMOylation), 27 SNO 1.0 (S-nitrosylation), 28 YNO21.0 (tyrosine nitration), 273 CCD1.0 (calpain cleavage), 36 Polo1.0 (polo-like kinases), 34 PUP1.0 (pupylation), 20 TSP1.0 (tyrosine sulfation), 25 PAIL2.0 (lysine acetylation), and 41 Lipid1.0 (lipidation) predicted modifications to SARS-CoV-2 viral proteins (Figure 5A) with data available at doi.org/10.6084/m9.figshare.12298790.v1 or within our VIStEDD tools (https://prokoplab.com/sars-cov-2/). Few modification predictions occur at highly conserved amino acids (Figure 5B) or within 5 amino acid conserved motifs (Figures 5C) based on our evolutionary analysis. Moreover, there are also very few unique modification predictions to the SARS-CoV-2 sequence not found throughout our evolutionary conservation. We have identified 22 modification predictions that are highly conserved and 28 sites poorly conserved including multiple phosphorylation, lipidation, and acetylation events (Figure 5D-F). The most highly conserved motif predicted modified is S816 of Spike (Figure 5G), where the amino acid is found surface exposed on a loop of the protein (Figure 5H-I). Future work is desperately needed to further refine the modification sites within SARS-CoV-2.

### SARS-CoV-2 Spike (S) interaction with host ACE2, SLC6A19, TMPRSS2 genomics

The most highly researched protein of SARS-CoV-2, the Spike (S) surface glycoprotein, is of most interest as it is the only portion outside of the virus that could be recognizable by B and T cells. The coronavirus spike protein (S) is the primary determinant of viral tropism and is responsible for receptor binding and membrane fusion. It is a large (approx.180 kDa) glycoprotein that is present on the viral surface as a prominent trimer, and it is composed of two domains, S1 and S2 (Belouzard et al., 2012). The S protein interacts with ACE2 to enter in cells, forming contacts with the ACE2-SLC6A19 dimer complex (Figure 6A). From 236 species of ACE2, we determined the conservation of ACE2 amino acids, suggesting poor conservation of the S-ACE2 contact across all of vertebrate evolution (Figure 6B). The lack of conservation at this site also suggests that ACE2 does not likely have a conserved interaction with another human protein that would compete with Spike for function. Structural mapping of human variants from 141,456 people of the gnomADv2 database for ACE2 suggest a few possible variants at the interface of S-ACE2 interaction (Figure 6C). To go from qualitative mapping to quantitative insights of human variants we utilized mds of S-ACE2-SLC6A19 complex, determining amino acids that correlate in movement between the proteins (Figure 6D). From these correlations we calculated the amino acids contributing to S-ACE interaction (red Figure 6E, Figure 6F), ACE2 dimerization (blue Figure 6E, Figure 6G), and ACE2-SLC6A19 interaction (magenta/yellow Figure 6E, Figure 6H-I).

**Figure 6.**
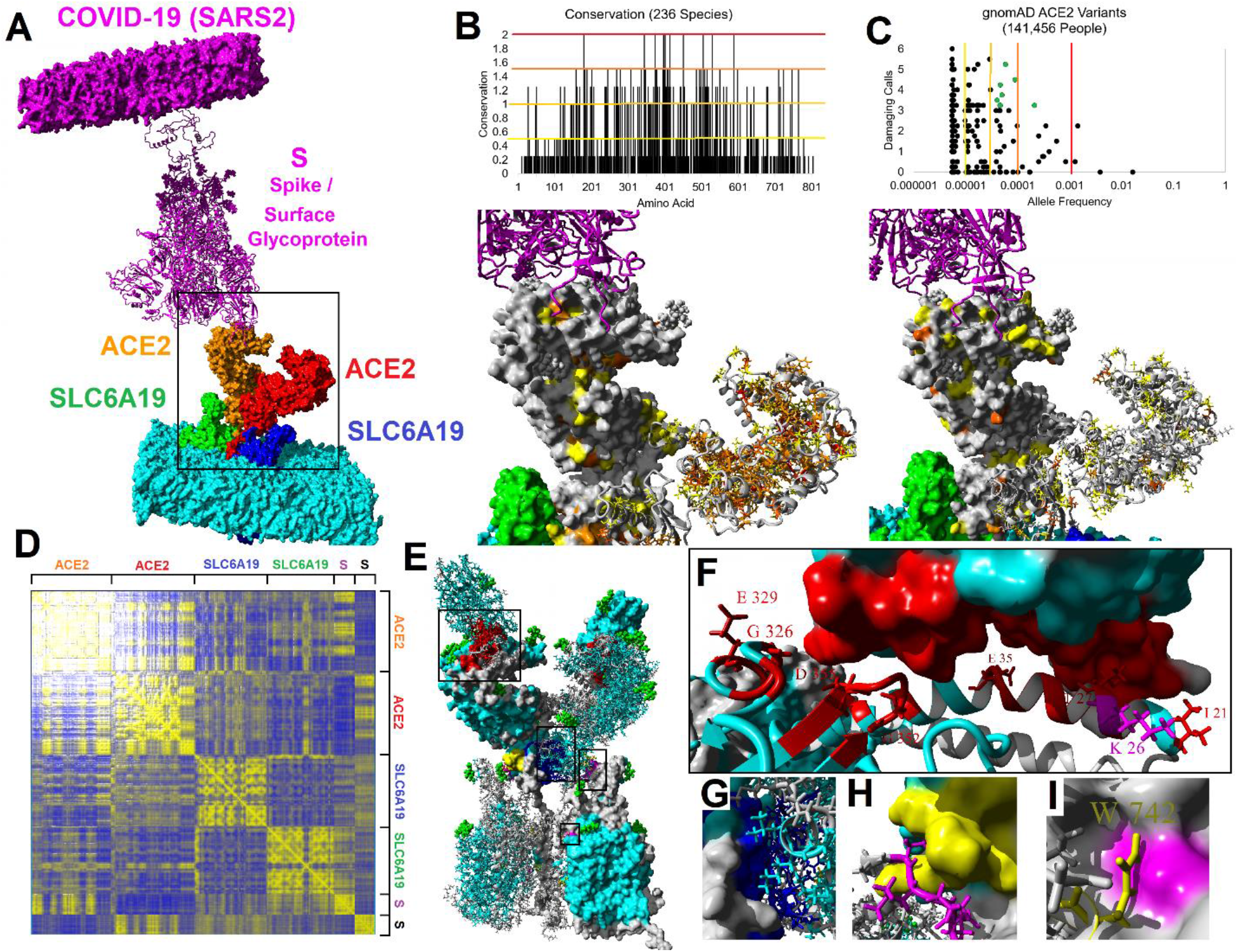
Spike-ACE2-SLC6A19 dynamics/evolution to human variants. **A)** Structural model of the Spike (magenta), ACE2 (red/orange), SLC6A19 (green/blue) in lipid membranes. Black box is region zoomed in in panels B-C. **B)** Conserved amino acids in 236 species ACE2. Color cutoffs: red=2 (highest), dark orange =1.5, light orange=1&1.25, yellow=1&0.5. **C)** Missense variants from 141,456 people for ACE2. Cutoffs for allele frequency are shown on the model. Several yellow low frequency variants can be seen at ACE2/Spike contact. **D)** Dynamics correlation matrix of interacting sites throughout simulation of the dimer complex. High correlations are shown in yellow. **E)** Amino acids in red are those that correlate in panel D between S and ACE2, those shown as side chains and labeled have human variants. The four boxes are the sites for panels F-I. **F)** Zoom in view of variants in ACE2 predicted to alter Spike contact (red). K26 is labeled in magenta with its uncertain but relatively common variant. **G)** Contact sites for ACE2 dimerization (blue). **H-I)** Contact sites between SLC6A19 (magenta) and ACE2 (yellow) at two different regions.

From this mds data along, with functional variant prediction tools (PolyPhen2, Provean, SIFT, Align-GVGD, and our conservation analysis), we systematically assessed functional human variants for ACE2, SLC6A19, and TMPRSS2. TMPRSS2 is involved in cleaving the complex for internalization (Hoffmann et al., 2020). Of these three proteins, ACE2 is the only one found on a sex chromosome (X-chromosome), linking it to male hemizygous status that elevates the impact of genomic variants. Variants included are linked to protein function (Table 2, Inclusion group = Functional), S contact (Table 2, Inclusion group = Spike Contact), SLC6A19-ACE2 contact (Table 2, Inclusion group = ACE2 Contact), posttranslational modifications (Table 2, Inclusion group = Glycosylation, Disulfide bond, or Phosphoserine), or known active site amino acids (Table 2, Inclusion group = Zinc Binding, Active Site).

**Table 2.** Top functional genomic variants of ACE2, SLC6A19, and TMPRSS2. The inclusion group consists of protein contacts based on molecular dynamic simulation correlations, protein modifications based on UniProt, and functional predictions. Damaging calls are a max of 6 based on conservation, PolyPhen2, Provean, SIFT, and Align-GVGD. Hemizygote count, max allele freq, and population are from gnomADv2.

| Inclusion | Protein | Chromosome | rsID | AA | Damaging Calls | Hemizygote Count | Max Allele Freq | Max Population |
| --- | --- | --- | --- | --- | --- | --- | --- | --- |
| Spike Contact | ACE2 | X | rs4646116 | K26R | 0 | 282 | 0.011876795 | Ashkenazi Jewish |
| Functional | SLC6A19 | 5 | rs781039193 | W530C | 5 | 0 | 0.000868709 | Latino |
| ACE2 Contact | SLC6A19 | 5 | rs374559483 | P351S | 1.5 | 0 | 0.000640872 | African |
| Functional | SLC6A19 | 5 | rs374527866 | R513S | 4.5 | 0 | 0.000603136 | East Asian |
| Functional | SLC6A19 | 5 | rs141487939 | T180M | 5.5 | 0 | 0.000601878 | African |
| Functional | SLC6A19 | 5 | rs200783817 | P73L | 5.25 | 0 | 0.000451309 | East Asian |
| Functional | SLC6A19 | 5 | rs752378013 | P109L | 5.5 | 0 | 0.000437158 | East Asian |
| Functional | SLC6A19 | 5 | rs141497538 | P230S | 5.5 | 0 | 0.000417246 | Other |
| Functional | SLC6A19 | 5 | rs552867213 | R328H | 5 | 0 | 0.000401091 | European (Finnish) |
| Glycosylation | ACE2 | X | rs761944150 | N546D | 2.5 | 1 | 0.00036784 | African |
| Functional | SLC6A19 | 5 | rs374976872 | A271V | 5.5 | 0 | 0.000300933 | East Asian |
| Functional | SLC6A19 | 5 | rs199795977 | A329T | 5 | 0 | 0.000289519 | Ashkenazi Jewish |
| Functional | SLC6A19 | 5 | rs1253340252 | Y209S | 4 | 0 | 0.000278242 | Other |
| Functional | SLC6A19 | 5 | rs139182948 | R95W | 5 | 0 | 0.000277701 | Other |
| Functional | SLC6A19 | 5 | rs369804798 | S314L | 5.5 | 0 | 0.000276932 | Other |
| Functional | SLC6A19 | 5 | rs142164435 | R328C | 5.5 | 0 | 0.000247927 | European (non-Finnish) |
| Functional | SLC6A19 | 5 | rs762989809 | R57C | 5.25 | 0 | 0.000218126 | East Asian |
| Glycosylation | ACE2 | X | rs143158922 | N103H | 0.5 | 0 | 0.000210615 | African |
| Zinc Binding | ACE2 | X | rs142984500 | H378R | 4.5 | 6 | 0.000194968 | European (non-Finnish) |
| Spike Contact | ACE2 | X | rs143936283 | E329G | 0 | 1 | 0.000190476 | Other |
| Functional | SLC6A19 | 5 | rs1403845937 | W56R | 5 | 0 | 0.000164582 | Other |
| Functional | SLC6A19 | 5 | rs201936518 | R95Q | 5 | 0 | 0.00016372 | Other |
| Functional | SLC6A19 | 5 | rs375879452 | T256M | 5.5 | 0 | 0.000150708 | East Asian |
| Functional | TMPRSS2 | 21 | rs762844469 | G391E | 5.25 | 0 | 0.000144601 | Latino |
| Spike Contact | ACE2 | X | rs1348114695 | E35K | 0 | 1 | 0.000144436 | East Asian |
| Functional | SLC6A19 | 5 | rs757679627 | G93R | 5.5 | 0 | 0.00013885 | Other |
| Functional | TMPRSS2 | 21 | rs1306483136 | S460R | 5.25 | 0 | 0.000138351 | Other |
| Functional | SLC6A19 | 5 | rs202220597 | T243M | 5.5 | 0 | 0.000130693 | South Asian |
| Functional | SLC6A19 | 5 | rs200745023 | L242P | 5 | 0 | 0.000130685 | South Asian |
| Functional | SLC6A19 | 5 | rs778015723 | P149L | 5.5 | 0 | 0.000130651 | South Asian |
| Functional | ACE2 | X | rs200745906 | P263S | 5.25 | 0 | 0.000125798 | European (non-Finnish) |
| Functional | SLC6A19 | 5 | rs765501634 | E405K | 5.25 | 0 | 0.000120318 | African |
| Functional | SLC6A19 | 5 | rs748703513 | T330I | 5 | 0 | 0.000114404 | European (non-Finnish) |
| Active Site | ACE2 | X | rs1395782023 | E375D | 3.25 | 1 | 0.000109585 | Latino |
| Functional | SLC6A19 | 5 | rs769457402 | E165K | 5.25 | 0 | 0.000108755 | East Asian |
| Functional | TMPRSS2 | 21 | rs145171279 | T459I | 5.5 | 0 | 0.000108731 | East Asian |
| Functional | SLC6A19 | 5 | rs756920378 | I325T | 5.25 | 0 | 0.000108731 | East Asian |
| Glycosylation | TMPRSS2 | 21 | rs1326192818 | N213K | 1.5 | 0 | 8.67553E-05 | Latino |
| Spike Contact | ACE2 | X | rs781255386 | T27A | 0 | 0 | 7.30327E-05 | Latino |
| Glycosylation | SLC6A19 | 5 | rs766784542 | N158T | 2.5 | 0 | 6.53381E-05 | South Asian |
| Functional | ACE2 | X | rs766319182 | M270V | 4 | 2 | 6.44787E-05 | European (non-Finnish) |
| Disulfide bond | TMPRSS2 | 21 | rs906113408 | C297Y | 6 | 0 | 5.99988E-05 | Latino |
| Glycosylation | SLC6A19 | 5 | rs146176472 | N258S | 1.5 | 0 | 4.06207E-05 | African |
| Functional | ACE2 | X | rs756358940 | I291K | 5.25 | 2 | 3.75756E-05 | European (non-Finnish) |
| Phosphoserine | SLC6A19 | 5 | rs754779609 | S17C | 2 | 0 | 3.57373E-05 | European (non-Finnish) |
| Spike Contact | ACE2 | X | rs961360700 | D355N | 4.25 | 0 | 2.59141E-05 | European (non-Finnish) |
| Spike Contact | ACE2 | X | rs778030746 | I21V | 0 | 2 | 2.44636E-05 | European (non-Finnish) |

47 variants are ranked by the max allele frequency within the subpopulations of gnomAD. The ACE2 variant K26R has the highest allele frequency of any variant within the table, but the conserved polar basic amino acid at the Spike contact likely does not impact binding. This means that there are only ultra-rare variants in these proteins, with SLC6A19 having the highest impact variants at 29, ACE2 with 13, and TMPRSS2 with 5. Outside of European non-Finnish population, the East Asian population carries 10 of these variants, “Other” (those individuals not falling into other populations) with 9, African with 6, Latino with 6, and South Asian with 4. A total of 31 of the variants are predicted with a score ≥4 (out of 6 max) to be functional variants, 6 at the Spike contact of ACE2, and 5 that would alter a glycosylation signal. The most interesting ACE2 variant, H378R (rs142984500), is found in 0.019% of European non-Finnish individuals, and has been observed as hemizygous in 6 males of gnomADv2, is one of the critical residues of the Zn binding that drives the enzymes function, and has not been previously published.

ACE2 hemizygous nature warrants an investigation of noncoding variants that might influence expression. The ACE2 gene contains a 5’ region that suggests most gene regulation for ACE2 to occur within this region (chrX:15,612,899-15,641,393, hg19). We extracted all noncoding variants followed by an assessment with RegulomeDB (Table S2). Five total variants are identified to potentially alter transcriptional regulation by RegulomeDB score. Two of these variants (rs4646118 and rs143185769) are found in ~9% of African individuals with hundreds of male hemizygotes identified within gnomADv3 whole genome data. This supports potential for noncoding variants of ACE2, with a higher allele frequency than that of coding variants, as contributors to increased susceptibility and within at-risk populations (African and Male) to SARS-CoV-2 infection.

## Discussion

SARS-CoV-2 represents a generational challenge to science, racing the clock next to a global pandemic that kills ~2% of those infected. The need to understand the viral structure is urgent regarding therapeutic targets, repurposing compounds, understanding zoonotic spread, and identifying gene variant risk factors in the human host that interact with the pathogen to increase spread and pathogenicity. In future studies, investigations of known human PPI to SARS-CoV-2, similar to ACE2, can determine how genetic variations of host proteins impact the disease course and susceptibility to the virus. Further insight into the genetics could provide useful information into the susceptibility or prognosis of those exposed to or infected with SARS-CoV-2. In this paper we generated a SARS-CoV-2 structural dynamicome database, integrating structural/dynamic insights with viral evolution for 24 proteins coded by SARS-CoV-2. VIStEDD has elucidated insights that include potential druggable targets, educational material describing each protein, and human variants that may impact the viral life cycle.

We show here two highly dynamic protein regions that have high conservation, indicative of PPI sites critical to viral infection and spreading. The first is the largely understudied role of the nsp6 conserved amino acids that interact with ATPases of vesicle trafficking (Gordon et al., 2020). Evolutionary conservation throughout thousands of *Coronaviridae* sequences is found in two regions of the protein that are likely found near each other in 3D space. Yet, to date no protein structures have ever been solved of nsp6 and publicly shared, representing a challenge to the structural biology community. ATPases are required for both endocytic and exocytic portions of the viral infections (Palokangas et al., 1994) and integral in the release of viral RNA into the cell (Hinton et al., 2009). The conserved amino acids are found on surfaces exposed on the I-TASSER generated predicted structure of nsp6 including multiple charged or aromatics residues. The conserved sites of nsp6 and the ATPases they interact with can likely be therapeutically targeted (Gordon et al., 2020). We also show here that the Nucleocapsid (N) protein has several highly conserved amino acids that contribute to a multimer structural organization with surface exposed conserved amino acids and the N-terminal region of the protein that may reflect sites of targeting to alter the ribosomal control of the protein.

Secondly, this new database presents many opportunities for use in the educational space. VIStEDD was generated through a team partnership known as Characterizing our DNA Exceptions (CODE) with the intent of bringing the mds and evolutionary data to undergraduate students across the US. The data can be used by anyone as the full dataset is publicly available (drive.google.com/drive/folders/1dXBJpLo3bay1JQ9BckUsVcTViv6P0w1q?usp=sharing). From this data, we provide high resolution figures of conservation mapped, structural files that can be opened in either YASARA or PyMOL tools, and molecular videos of the molecules rotating with conservation. For the 3D files, we have provided a vrml file for each protein that can be fed to any 3D printer, with our file containing colors for conservation as well. To expedite 3D printing, we have provided all the vrml files to Shapeways to allow at cost printing of the proteins (Table S1), where the files can be used for education.

The biophysical and structural evidence suggested that SARS-CoV-2 may bind ACE2 with a much higher affinity than SARS-CoV (Wrapp et al., 2020). Our group had previously investigated the evolution of ACE2 throughout species including mapping variants within rat populations (Prokop et al., 2015), a model system for studying the Renin-Angiotensin Aldosterone system (known as the RAAS). We have expanded those tools here, generating ACE2 models for 235 species from mammals to birds/fish, where each of the models are energy minimized with the S protein interaction. That database can be used by groups to investigate species where SARS-CoV-2 may be able to enter cells, with variants that enhance or inhibit binding. With the S-ACE2-SLC6A19 complex resolved and mds available, a systematic quantitative map of human variants was created. Few functional variants were identified in ACE2, SLC6A19, or TMPRSS2. Moreover, none of the variants identified are common, all falling sub 1% of the global population. In SLC6A19 or TMPRSS2 these rare variants would have minimal outcomes as they would rarely, if ever, reach homozygosity to cause 100% of the proteins to be influenced by variants. However, ACE2 falls on the X-chromosome, linking it to male specific hemizygous influence. Several of the top rare ACE2 variants identified within this paper have been seen to have hemizygous variants, suggesting a dominant outcome. While functional missense variants in ACE2 are ultra-rare, noncoding variants reach slightly higher allele frequencies, namely rs4646118 and rs143185769. This would suggest that as genome sequencing of patients with SARS-CoV-2 occurs, we should focus on analysis of ultra-rare missense variants listed within this paper and more importantly on several likely functional noncoding variants.

Through the quantitative dissection of the SARS-CoV-2 encoded proteins and their interaction partners we have developed a database (VIStEDD) of information that can be used to advance our knowledge. From the ability to regulate these interactions with pharmaceutical intervention to understanding how host genomics can influence the viral biology. VIStEDD will allow for a more robust insight into SARS-CoV-2 biology.

## Methods

### Protein modeling and molecular dynamic simulations

Similar to our lab’s analysis of human variants, we have assessed the SARS-CoV-2 proteins using our previous established workflow (Prokop et al., 2017). Protein modeling was performed by utilizing YASARA homology modeling (Krieger et al., 2009; Krieger and Vriend, 2015) when a structural template was available that matched the sequences listed in Table 1. Homology modeling is the preferred platform as it allows molecules associated with the proteins to be included in the protein structure, including Zn ions critical to folding of Zn-fingers within the Papain-like proteinase, nsp10, RNA-directed RNA polymerase, Helicase, and Guanine-N7 methyltransferase. The transmembrane portions of S were manually cleaned and clustered allowing for insertion into a PEA membrane before molecular dynamic simulations (mds). For those proteins without structural homologs we utilized models that are part of the I-TASSER SARS-CoV-2 database (Zhang et al., 2020). Each of these models were then fed through homology modeling in YASARA to normalize energetic predictions to the homology models. Molecular dynamic simulations (mds) was performed on each of the proteins in YASARA (Krieger and Vriend, 2015) using the AMBER14 force field (Duan et al., 2003), 0.997g/mL explicit water, NaCl at 0.9 mass fraction, a pH of 7.4 for protonation predictions, saving trajectory files every 25 picoseconds for 20 nanoseconds total. The trajectory files were analyzed with the YASARA md_analyze and md_analyzeres macros, generating an HTML file present in each of the protein report folders. If this report folder is downloaded and the HTML file opened it generates a full report of the protein dynamics including multiple figures of analysis. Additionally, all the tab delineated analysis files are within the tab folder of VIStEDD proteins and the trajectory files allowing for reanalysis of trajectory.

### Database information generation

From the models and mds, we generated files for VIStEDD. Sequences (within the genomics folder of each protein) were extracted using the sequences listed in table 1 with BLASTp against the non-redundant protein sequences (nr) and aligned using ClustalW (Larkin et al., 2007). An amino acid matrix of all sequences was generated in MEGA (Tamura et al., 2011, p. 6) followed by calculating the percent of all amino acids at each spot that are the same as in SARS-CoV-2. The conservation of each protein was normalized using a z-score ([value-mean]/standard deviation) for comparison across all proteins. The generated models were loaded into YASARA (Z-score 0-0.5 = yellow, 0.5-1 = color 166, 1-1.5 = color 157, 1.5-2 = color 145, >2 = red) or PyMOL (Z-score 0-0.5 = yellow, 0.5-1 = bright orange, 1-1.5 = orange, 1.5-2 = warmpink, >2 = red), colored based on conservation, and saved as respective scene files for the tool. The YASARA colored molecule was saved as a Y-axis rotation in mpg and converted into mp4, which is more amenable to PowerPoint. Within PyMOL the structure was also exported as a vrml file for 3D printing. Motifs and posttranslational modification predictions for the N protein were generated using ELM (Dinkel et al., 2012) and NetPhos3.1 (Blom et al., 2004).

### Human variant analysis

Models for ACE2 and SLC6A19 were homology modeled using PDB 6M17 (S-ACE2-SLC6A19) followed by alignment back onto the complex. TMPRSS2 was homology modeled using YASARA followed by manual correction of the transmembrane helix. Each of the two models was embedded into a PEA lipid membrane using the YASARA macro md_runmembrane followed by mds as done on SARS-CoV-2 proteins. Vertebrate sequences of the three proteins were extracted using NCBI orthologs for the transcript, open reading frames assessed using Transdecoder (Haas et al., 2013), and aligned using ClustalW codons in MEGA. Conservation was performed on the data as previously published (Prokop et al., 2018). Genomic missense variants were extracted from gnomADv2 for each of the three genes followed by assessment using PolyPhen2 (Adzhubei et al., 2010), Provean (Choi and Chan, 2015), SIFT (Ng and Henikoff, 2003), and Align-GVGD (Tavtigian et al., 2006). The ACE2 regulatory region was identified using Roadmap Epigenomics 18-state model (Roadmap Epigenomics Consortium et al., 2015, p. 111) followed by the extraction of all gnomADv3 variants and assessment with RegulomeDB (Boyle et al., 2012).

## Acknowledgements

Funding for this study were from NIH R01-GM108618 (JAC) and K01-ES025435 (JWP), Michigan State University, and Spectrum Health.

## Supplemental File

**Table S1.**
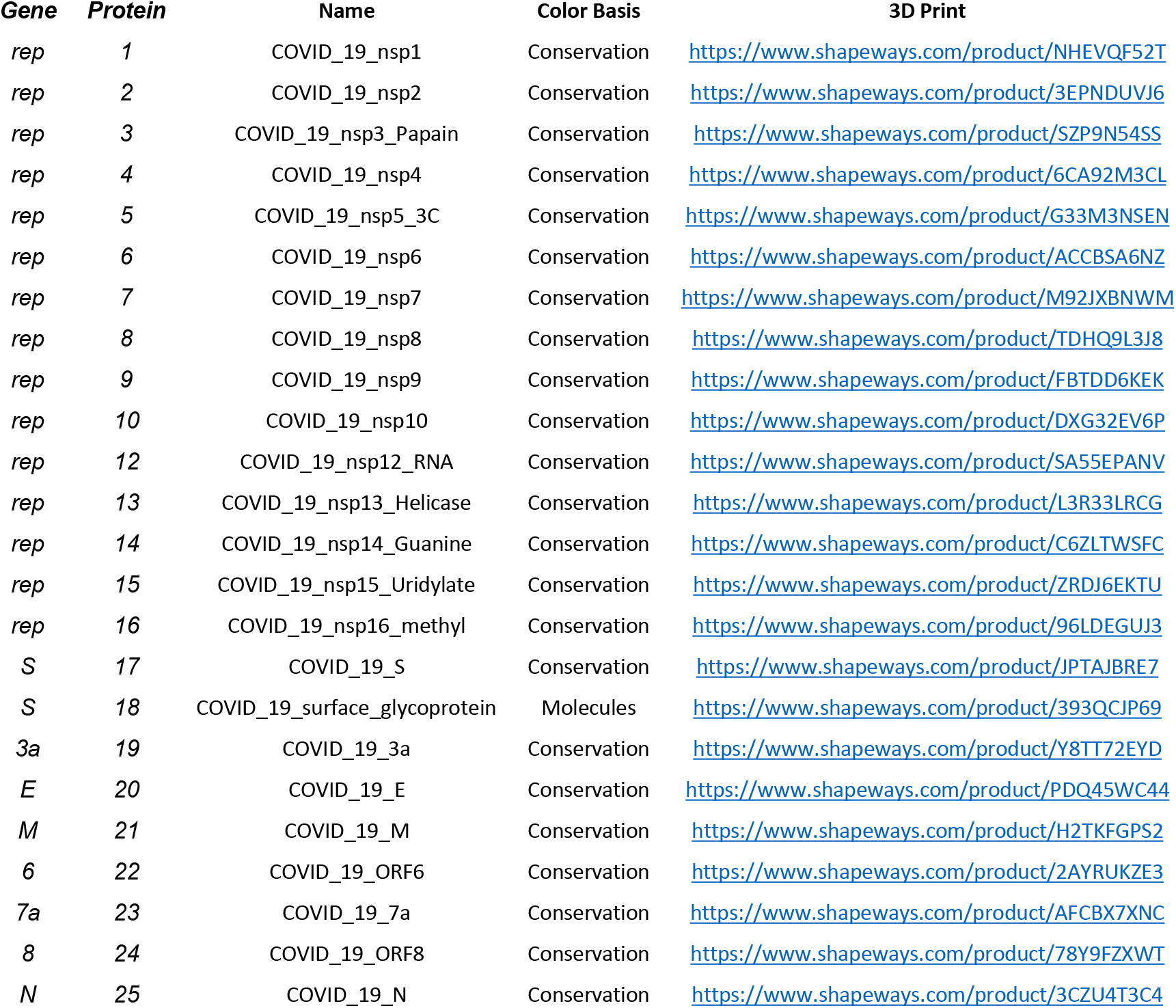
3D printing web links.

**Table S2.**
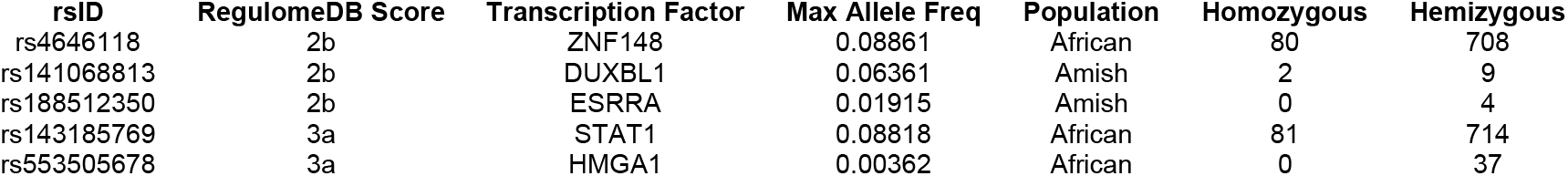
Top Noncoding variants of ACE2. Shown are the top five noncoding variants in the promoter of ACE2 with RegulomeDB score, predicted transcription factor altered binding, and gnomADv3 allele information.

## References

Adzhubei IA, Schmidt S, Peshkin L, Ramensky VE, Gerasimova A, Bork P, Kondrashov AS, Sunyaev SR. 2010. A method and server for predicting damaging missense mutations. Nat Methods 7:248–249. doi:10.1038/nmeth0410-248

Andersen KG, Rambaut A, Lipkin WI, Holmes EC, Garry RF. 2020. The proximal origin of SARS-CoV-2. Nature Medicine 26:450–452. doi:10.1038/s41591-020-0820-9

Angelini MM, Akhlaghpour M, Neuman BW, Buchmeier MJ. 2013. Severe Acute Respiratory Syndrome Coronavirus Nonstructural Proteins 3, 4, and 6 Induce Double-Membrane Vesicles. mBio 4. doi:10.1128/mBio.00524-13

Barkauskas CE, Cronce MJ, Rackley CR, Bowie EJ, Keene DR, Stripp BR, Randell SH, Noble PW, Hogan BLM. 2013. Type 2 alveolar cells are stem cells in adult lung. J Clin Invest 123:3025–3036. doi:10.1172/JCI68782

Belouzard S, Millet JK, Licitra BN, Whittaker GR. 2012. Mechanisms of coronavirus cell entry mediated by the viral spike protein. Viruses 4:1011–1033. doi:10.3390/v4061011

Blom N, Sicheritz-Pontén T, Gupta R, Gammeltoft S, Brunak S. 2004. Prediction of post-translational glycosylation and phosphorylation of proteins from the amino acid sequence. Proteomics 4:1633–1649. doi:10.1002/pmic.200300771

Boyle AP, Hong EL, Hariharan M, Cheng Y, Schaub MA, Kasowski M, Karczewski KJ, Park J, Hitz BC, Weng S, Cherry JM, Snyder M. 2012. Annotation of functional variation in personal genomes using RegulomeDB. Genome Res 22:1790–1797. doi:10.1101/gr.137323.112

Ceraolo C, Giorgi FM. 2020. Genomic variance of the 2019-nCoV coronavirus. J Med Virol 92:522–528. doi:10.1002/jmv.25700

Choi Y, Chan AP. 2015. PROVEAN web server: a tool to predict the functional effect of amino acid substitutions and indels. Bioinformatics 31:2745–2747. doi:10.1093/bioinformatics/btv195

Cottam EM, Whelband MC, Wileman T. 2014. Coronavirus NSP6 restricts autophagosome expansion. Autophagy 10:1426–1441. doi:10.4161/auto.29309

Dinkel H, Michael S, Weatheritt RJ, Davey NE, Van Roey K, Altenberg B, Toedt G, Uyar B, Seiler M, Budd A, Jödicke L, Dammert MA, Schroeter C, Hammer M, Schmidt T, Jehl P, McGuigan C, Dymecka M, Chica C, Luck K, Via A, Chatr-Aryamontri A, Haslam N, Grebnev G, Edwards RJ, Steinmetz MO, Meiselbach H, Diella F, Gibson TJ. 2012. ELM--the database of eukaryotic linear motifs. Nucleic Acids Res 40:D242–251. doi:10.1093/nar/gkr1064

Dong E, Du H, Gardner L. 2020. An interactive web-based dashboard to track COVID-19 in real time. Lancet Infect Dis. doi:10.1016/S1473-3099(20)30120-1

Duan Y, Wu C, Chowdhury S, Lee MC, Xiong G, Zhang W, Yang R, Cieplak P, Luo R, Lee T, Caldwell J, Wang J, Kollman P. 2003. A point-charge force field for molecular mechanics simulations of proteins based on condensed-phase quantum mechanical calculations. J Comput Chem 24:1999–2012. doi:10.1002/jcc.10349

Fan Z, Zhuo Y, Tan X, Zhou Z, Yuan J, Qiang B, Yan J, Peng X, Gao GF. 2006. SARS-CoV nucleocapsid protein binds to hUbc9, a ubiquitin conjugating enzyme of the sumoylation system. J Med Virol 78:1365–1373. doi:10.1002/jmv.20707

Fung TS, Liu DX. 2018. Post-translational modifications of coronavirus proteins: roles and function. Future Virol 13:405–430. doi:10.2217/fvl-2018-0008

Gordon DE, Jang GM, Bouhaddou M, Xu J, Obernier K, O’Meara MJ, Guo JZ, Swaney DL, Tummino TA, Huettenhain R, Kaake RM, Richards AL, Tutuncuoglu B, Foussard H, Batra J, Haas K, Modak M, Kim M, Haas P, Polacco BJ, Braberg H, Fabius JM, Eckhardt M, Soucheray M, Bennett MJ, Cakir M, McGregor MJ, Li Q, Naing ZZC, Zhou Y, Peng S, Kirby IT, Melnyk JE, Chorba JS, Lou K, Dai SA, Shen W, Shi Y, Zhang Z, Barrio-Hernandez I, Memon D, Hernandez-Armenta C, Mathy CJP, Perica T, Pilla KB, Ganesan SJ, Saltzberg DJ, Ramachandran R, Liu X, Rosenthal SB, Calviello L, Venkataramanan S, Liboy-Lugo J, Lin Y, Wankowicz SA, Bohn M, Sharp PP, Trenker R, Young JM, Cavero DA, Hiatt J, Roth TL, Rathore U, Subramanian A, Noack J, Hubert M, Roesch F, Vallet T, Meyer B, White KM, Miorin L, Rosenberg OS, Verba KA, Agard D, Ott M, Emerman M, Ruggero D, García-Sastre A, Jura N, Zastrow M von, Taunton J, Ashworth A, Schwartz O, Vignuzzi M, d’Enfert C, Mukherjee S, Jacobson M, Malik HS, Fujimori DG, Ideker T, Craik CS, Floor S, Fraser JS, Gross J, Sali A, Kortemme T, Beltrao P, Shokat K, Shoichet BK, Krogan NJ. 2020. A SARS-CoV-2-Human Protein-Protein Interaction Map Reveals Drug Targets and Potential Drug-Repurposing. bioRxiv 2020.03.22.002386. doi:10.1101/2020.03.22.002386

Guarner J. 2020. Three Emerging Coronaviruses in Two DecadesThe Story of SARS, MERS, and Now COVID-19. Am J Clin Pathol 153:420–421. doi:10.1093/ajcp/aqaa029

Haas BJ, Papanicolaou A, Yassour M, Grabherr M, Blood PD, Bowden J, Couger MB, Eccles D, Li B, Lieber M, MacManes MD, Ott M, Orvis J, Pochet N, Strozzi F, Weeks N, Westerman R, William T, Dewey CN, Henschel R, LeDuc RD, Friedman N, Regev A. 2013. De novo transcript sequence reconstruction from RNA-seq using the Trinity platform for reference generation and analysis. Nat Protoc 8:1494–1512. doi:10.1038/nprot.2013.084

Hamming I, Timens W, Bulthuis MLC, Lely AT, Navis GJ, van Goor H. 2004. Tissue distribution of ACE2 protein, the functional receptor for SARS coronavirus. A first step in understanding SARS pathogenesis. J Pathol 203:631–637. doi:10.1002/path.1570

Hinton A, Bond S, Forgac M. 2009. V-ATPase functions in normal and disease processes. Pflugers Arch 457:589–598. doi:10.1007/s00424-007-0382-4

Hoffmann M, Kleine-Weber H, Schroeder S, Krüger N, Herrler T, Erichsen S, Schiergens TS, Herrler G, Wu N-H, Nitsche A, Müller MA, Drosten C, Pöhlmann S. 2020. SARS-CoV-2 Cell Entry Depends on ACE2 and TMPRSS2 and Is Blocked by a Clinically Proven Protease Inhibitor. Cell. doi:10.1016/j.cell.2020.02.052

Kasmi Y, Khataby K, Souiri A, Ennaji MM. 2020. Chapter 7 - Coronaviridae: 100,000 Years of Emergence and Reemergence In: Ennaji MM, editor. Emerging and Reemerging Viral Pathogens. Academic Press. pp. 127–149. doi:10.1016/B978-0-12-819400-3.00007-7

Krieger E, Joo K, Lee Jinwoo, Lee Jooyoung, Raman S, Thompson J, Tyka M, Baker D, Karplus K. 2009. Improving physical realism, stereochemistry, and side-chain accuracy in homology modeling: Four approaches that performed well in CASP8. Proteins 77 Suppl 9:114–122. doi:10.1002/prot.22570

Krieger E, Vriend G. 2015. New ways to boost molecular dynamics simulations. J Comput Chem 36:996–1007. doi:10.1002/jcc.23899

Kuba K, Imai Y, Rao S, Gao H, Guo F, Guan B, Huan Y, Yang P, Zhang Y, Deng W, Bao L, Zhang B, Liu G, Wang Z, Chappell M, Liu Y, Zheng D, Leibbrandt A, Wada T, Slutsky AS, Liu D, Qin C, Jiang C, Penninger JM. 2005. A crucial role of angiotensin converting enzyme 2 (ACE2) in SARS coronavirus|[ndash]|induced lung injury. Nature Medicine 11:875–879. doi:10.1038/nm1267

Larkin MA, Blackshields G, Brown NP, Chenna R, McGettigan PA, McWilliam H, Valentin F, Wallace IM, Wilm A, Lopez R, Thompson JD, Gibson TJ, Higgins DG. 2007. Clustal W and Clustal X version 2.0. Bioinformatics 23:2947–2948. doi:10.1093/bioinformatics/btm404

Malle L. 2020. A map of SARS-CoV-2 and host cell interactions. Nature Reviews Immunology 1–1. doi:10.1038/s41577-020-0318-1

Mehta P, McAuley DF, Brown M, Sanchez E, Tattersall RS, Manson JJ, HLH Across Speciality Collaboration, UK. 2020. COVID-19: consider cytokine storm syndromes and immunosuppression. Lancet 395:1033–1034. doi:10.1016/S0140-6736(20)30628-0

Nal B, Chan C, Kien F, Siu L, Tse J, Chu K, Kam J, Staropoli I, Crescenzo-Chaigne B, Escriou N, van der Werf S, Yuen K-Y, Altmeyer R. 2005. Differential maturation and subcellular localization of severe acute respiratory syndrome coronavirus surface proteins S, M and E. J Gen Virol 86:1423–1434. doi:10.1099/vir.0.80671-0

Ng PC, Henikoff S. 2003. SIFT: Predicting amino acid changes that affect protein function. Nucleic Acids Res 31:3812–3814.

Oostra M, de Haan C a. M, de Groot RJ, Rottier PJM. 2006. Glycosylation of the severe acute respiratory syndrome coronavirus triple-spanning membrane proteins 3a and M. J Virol 80:2326–2336. doi:10.1128/JVI.80.5.2326-2336.2006

Palokangas H, Metsikkö K, Väänänen K. 1994. Active vacuolar H+ATPase is required for both endocytic and exocytic processes during viral infection of BHK-21 cells. J Biol Chem 269:17577–17585.

Prokop JW, Lazar J, Crapitto G, Smith DC, Worthey EA, Jacob HJ. 2017. Molecular modeling in the age of clinical genomics, the enterprise of the next generation. J Mol Model 23:75. doi:10.1007/s00894-017-3258-3

Prokop JW, Petri V, Shimoyama ME, Watanabe IKM, Casarini DE, Leeper TC, Bilinovich SM, Jacob HJ, Santos RAS, Martins AS, Araujo FC, Reis FM, Milsted A. 2015. Structural libraries of protein models for multiple species to understand evolution of the renin-angiotensin system. Gen Comp Endocrinol 215:106–116. doi:10.1016/j.ygcen.2014.09.010

Prokop JW, Yeo NC, Ottmann C, Chhetri SB, Florus KL, Ross EJ, Sosonkina N, Link BA, Freedman BI, Coppola CJ, McDermott-Roe C, Leysen S, Milroy L-G, Meijer FA, Geurts AM, Rauscher FJ, Ramaker R, Flister MJ, Jacob HJ, Mendenhall EM, Lazar J. 2018. Characterization of Coding/Noncoding Variants forSHROOM3in Patients with CKD. J Am Soc Nephrol. doi:10.1681/ASN.2017080856

Prompetchara E, Ketloy C, Palaga T. 2020. Immune responses in COVID-19 and potential vaccines: Lessons learned from SARS and MERS epidemic. Asian Pac J Allergy Immunol 38:1–9. doi:10.12932/AP-200220-0772

Roadmap Epigenomics Consortium, Kundaje A, Meuleman W, Ernst J, Bilenky M, Yen A, Heravi-Moussavi A, Kheradpour P, Zhang Z, Wang J, Ziller MJ, Amin V, Whitaker JW, Schultz MD, Ward LD, Sarkar A, Quon G, Sandstrom RS, Eaton ML, Wu Y-C, Pfenning AR, Wang X, Claussnitzer M, Liu Y, Coarfa C, Harris RA, Shoresh N, Epstein CB, Gjoneska E, Leung D, Xie W, Hawkins RD, Lister R, Hong C, Gascard P, Mungall AJ, Moore R, Chuah E, Tam A, Canfield TK, Hansen RS, Kaul R, Sabo PJ, Bansal MS, Carles A, Dixon JR, Farh K-H, Feizi S, Karlic R, Kim A-R, Kulkarni A, Li D, Lowdon R, Elliott G, Mercer TR, Neph SJ, Onuchic V, Polak P, Rajagopal N, Ray P, Sallari RC, Siebenthall KT, Sinnott-Armstrong NA, Stevens M, Thurman RE, Wu J, Zhang B, Zhou X, Beaudet AE, Boyer LA, De Jager PL, Farnham PJ, Fisher SJ, Haussler D, Jones SJM, Li W, Marra MA, McManus MT, Sunyaev S, Thomson JA, Tlsty TD, Tsai L-H, Wang W, Waterland RA, Zhang MQ, Chadwick LH, Bernstein BE, Costello JF, Ecker JR, Hirst M, Meissner A, Milosavljevic A, Ren B, Stamatoyannopoulos JA, Wang T, Kellis M. 2015. Integrative analysis of 111 reference human epigenomes. Nature 518:317–330. doi:10.1038/nature14248

Shi S, Qin M, Shen B, Cai Y, Liu T, Yang F, Gong W, Liu X, Liang J, Zhao Q, Huang H, Yang B, Huang C. 2020. Association of Cardiac Injury With Mortality in Hospitalized Patients With COVID-19 in Wuhan, China. JAMA Cardiol. doi:10.1001/jamacardio.2020.0950

Sungnak W, Huang N, Bécavin C, Berg M, Queen R, Litvinukova M, Talavera-López C, Maatz H, Reichart D, Sampaziotis F, Worlock KB, Yoshida M, Barnes JL, HCA Lung Biological Network. 2020. SARS-CoV-2 entry factors are highly expressed in nasal epithelial cells together with innate immune genes. Nat Med. doi:10.1038/s41591-020-0868-6

Surjit M, Kumar R, Mishra RN, Reddy MK, Chow VTK, Lal SK. 2005. The severe acute respiratory syndrome coronavirus nucleocapsid protein is phosphorylated and localizes in the cytoplasm by 14-3-3-mediated translocation. J Virol 79:11476–11486. doi:10.1128/JVI.79.17.11476-11486.2005

Surjit M, Lal SK. 2008. The SARS-CoV nucleocapsid protein: a protein with multifarious activities. Infect Genet Evol 8:397–405. doi:10.1016/j.meegid.2007.07.004

Tamura K, Peterson D, Peterson N, Stecher G, Nei M, Kumar S. 2011. MEGA5: molecular evolutionary genetics analysis using maximum likelihood, evolutionary distance, and maximum parsimony methods. Mol Biol Evol 28:2731–2739. doi:10.1093/molbev/msr121

Tavtigian SV, Deffenbaugh AM, Yin L, Judkins T, Scholl T, Samollow PB, de Silva D, Zharkikh A, Thomas A. 2006. Comprehensive statistical study of 452 BRCA1 missense substitutions with classification of eight recurrent substitutions as neutral. J Med Genet 43:295–305. doi:10.1136/jmg.2005.033878

Uhal BD, Dang M, Dang V, Llatos R, Cano E, Abdul-Hafez A, Markey J, Piasecki CC, Molina-Molina M. 2013. Cell cycle dependence of ACE-2 explains downregulation in idiopathic pulmonary fibrosis. European Respiratory Journal 42:198–210. doi:10.1183/09031936.00015612

Wada M, Lokugamage KG, Nakagawa K, Narayanan K, Makino S. 2018. Interplay between coronavirus, a cytoplasmic RNA virus, and nonsense-mediated mRNA decay pathway. PNAS 115:E10157–E10166. doi:10.1073/pnas.1811675115

Walls AC, Park Y-J, Tortorici MA, Wall A, McGuire AT, Veesler D. 2020. Structure, Function, and Antigenicity of the SARS-CoV-2 Spike Glycoprotein. Cell 181:281–292.e6. doi:10.1016/j.cell.2020.02.058

Wrapp D, Wang N, Corbett KS, Goldsmith JA, Hsieh C-L, Abiona O, Graham BS, McLellan JS. 2020. Cryo-EM structure of the 2019-nCoV spike in the prefusion conformation. Science 367:1260–1263. doi:10.1126/science.abb2507

Wu C, Liu Y, Yang Y, Zhang P, Zhong W, Wang Y, Wang Q, Xu Y, Li M, Li X, Zheng M, Chen L, Li H. 2020. Analysis of therapeutic targets for SARS-CoV-2 and discovery of potential drugs by computational methods. Acta Pharmaceutica Sinica B. doi:10.1016/j.apsb.2020.02.008

Xiao X, Chakraborti S, Dimitrov AS, Gramatikoff K, Dimitrov DS. 2003. The SARS-CoV S glycoprotein: expression and functional characterization. Biochem Biophys Res Commun 312:1159–1164. doi:10.1016/j.bbrc.2003.11.054

Yan R, Zhang Y, Li Y, Xia L, Guo Y, Zhou Q. 2020. Structural basis for the recognition of SARS-CoV-2 by full-length human ACE2. Science 367:1444–1448. doi:10.1126/science.abb2762

Zhang C, Zheng W, Huang X, Bell EW, Zhou X, Zhang Y. 2020. Protein Structure and Sequence Reanalysis of 2019-nCoV Genome Refutes Snakes as Its Intermediate Host and the Unique Similarity between Its Spike Protein Insertions and HIV-1. J Proteome Res 19:1351–1360. doi:10.1021/acs.jproteome.0c00129

Zhou F, Yu T, Du R, Fan G, Liu Y, Liu Z, Xiang J, Wang Y, Song B, Gu X, Guan L, Wei Y, Li H, Wu X, Xu J, Tu S, Zhang Y, Chen H, Cao B. 2020. Clinical course and risk factors for mortality of adult inpatients with COVID-19 in Wuhan, China: a retrospective cohort study. Lancet 395:1054–1062. doi:10.1016/S0140-6736(20)30566-3

